# Glycosaminoglycans act as activators of peptidylarginine deiminase 4

**DOI:** 10.1101/2024.06.17.599283

**Authors:** Grzegorz P. Bereta, Ewa Bielecka, Karolina Marzec, Łukasz Pijanowski, Artur Biela, Piotr Wilk, Marta Kamińska, Jakub Nowak, Elżbieta Wątor, Przemysław Grudnik, Dominik Kowalczyk, Joanna Kozieł, Piotr Mydel, Marcin Poręba, Tomasz Kantyka

## Abstract

Peptidylarginine deiminase 4 (PAD4) is a citrullinating enzyme that is gathering increasing attention due to its possible involvement in physiological processes as well as in the pathogenesis of diseases like rheumatoid arthritis or thrombosis. PAD4 is activated by calcium ions, but the details of this mechanism are elusive, because in the human body, Ca^2+^ concentrations are too low for full activity. Given that glycosaminoglycans (GAGs) are also implicated in the development and progression of rheumatoid arthritis, we investigated the activation of PAD4 by GAGs using heparin as a model. We employed activity assays, chromatography techniques, molecular interaction measurements (MST and SPR), FACS, and immunocytochemistry to demonstrate the activation of PAD4 by GAGs. Our data show that PAD4 binds heparin with high affinity and forms high molecular weight complexes with heparin, consistent with heparin-bound tetramer formation. Heparin activates PAD4 by increasing the enzyme’s Ca^2+^ affinity threefold. We also show that the effectiveness of activation with heparin depends on the length of GAG used and its negative charge. Direct measurement of heparin binding to PAD4 confirmed tight interaction with nanomolar affinity. Mutagenesis of regions likely responsible for heparin binding showed that dimerization of PAD4 is necessary for efficient activation, but the distinct binding site was not determined as interaction with heparin likely occurs over larger surface of PAD4. Furthermore, we show that other GAG family members, including heparan and chondroitin sulphates, are also able to activate PAD4. We also found that disturbed production of GAGs by CHO cells results in reduced PAD4 binding efficiency. Finally, heparin induces NETosis in hPMNs in concentration-dependent manner, as measured by the release of DNA and citrullination of histone H3. In summary, we identify the first natural coactivator of PAD4, which is present in all individuals, potentially explaining the regulation of PAD4 activity in physiological conditions, and providing new insight into the development of rheumatoid arthritis and other PAD4-related diseases.

## Main Text

Posttranslational modifications of proteins play important roles in both physiological functions and pathological conditions. Citrullination of arginine residues, which results in the loss of positive charge on the side chain, is performed by PAD enzymes, and PAD4 is one of ubiquitous members of this family (*1*). PAD4 is associated with rheumatoid arthritis (RA), where citrullinated epitopes are the main pathological factor (*2*–*4*). The activity of PAD4 is crucial for the process of NETosis, as it is necessary for chromatin decondensation through histone citrullination (*5*–*8*). This is made possible due to the nuclear localization signal identified in PAD4, which is unique among human PAD enzymes (*9*). The active form of PAD4 exists as homodimers, and dimerization greatly impacts enzyme activity, as monomeric mutants have been observed to be less active (*10*–*12*). The activity of this enzyme also depends on the concentration of calcium ions; a monomer of PAD4 has five binding sites for Ca^2+^, and their saturation results in conformational changes leading to maturation of the active site (*13*–*15*). Half-maximal activation of PAD4 is achieved at calcium concentrations between 300 to 600 µM in biochemical assays (*14, 15*), while concentration above 5 mM are required for full activity. The physiological concentration of free calcium reaches around 1 mM in the blood, while being orders of magnitude lower in the resting cells (*16, 17*). Cytoplasmic and nuclear PAD4 are activated by calcium influx, yet free calcium concentrations are still significantly below 1 mM under these conditions. In neutrophil-like cells, PAD4 citrullinates not only RA epitopes, but also multiple other proteins, including regulatory ones (*18*). It has been shown that immunization of susceptible mice with PAD4 led to the development of antibodies against citrullinated proteins, and PAD4/2 were found to be active in synovial fluid of RA patients, suggesting that PAD4 may act extracellularly (*19, 20*). Blood glycosaminoglycan (GAG) levels have been correlated with RA severity in humans, and GAG injections induced RA-like symptoms in mouse model, suggesting GAG contribution to RA onset (*21, 22*). Additionally, heparin, a common GAG molecule, was found to directly stimulate NETosis (*23*). Connecting GAGs with RA initiation and progression, heparin’s ability to stimulate PAD4-dependent NETosis, and the apparent inability of PAD4 to become fully active under physiological calcium concentrations, we hypothesized that GAGs, exemplified by heparin can directly modulate PAD4 activity, allowing efficient activation at sub-optimal Ca^2+^ concentrations.

### Heparin binds and activates PAD4

Heparin chromatographic resins are commonly used as universal ion-exchange or specific affinity media for proteins. We therefore tested the binding of recombinant human PAD4 to an analytical heparin column (Fig. 1A) and found that the protein eluted around 1.4 M salt concentration, later than expected for a simple ion-exchange interaction, suggesting stronger specific binding. We verified this observation by size exclusion chromatography (SEC) for PAD4 alone and in the presence of heparin. In the PAD4 elution profile, we observed peaks corresponding to monomer, containing majority of protein, together with dimer and a larger form (possibly tetramer). After the addition of heparin, the majority of PAD4 eluted as a novel complex, larger than any form of PAD4 alone, while the monomeric form was detected in small quantities (Fig. 1B). The estimated molecular weight of this novel, heparin-induced complex is consistent with the formation of the GAG-bound tetramer of PAD4. To investigate the potential activity of this novel PAD4-heparin complex, we measured enzymatic activity by HPLC using fluorescently labelled di-or tripeptide substrates in the presence of sub-activatory Ca^2+^ concentration (0.1 mM). Heparin activated PAD4 with an apparent K_D_ below 1 nM (Fig. 1C) with both substrates. We verified the potential activation mechanism by testing PAD4 calcium dependence in the presence of an activating heparin concentration (1 µM), and observed a decrease in half-activatory calcium concentration Ca_0.5_ on both peptide substrates from 155.8 – 148.2 µM to 77.4 – 55.1 µM without and with heparin, respectively (Fig. 1D). We further validated these results using an established PAD4 activity assay with a colorimetric, small molecular substrate BAEE and measured a heparin apparent K_D_ of 15.3 nM and Ca_0.5_ of 92.9 µM in the presence of heparin, contrasted by 367 µM without it (Fig. 1E, F). We directly measured the PAD4 – heparin interaction using microscale thermophoresis in two alternate settings: measuring the signal from FITC-labelled heparin with unlabeled PAD4, or from intrinsic PAD4 fluorescence with unlabeled heparin, obtaining K_D_ around 8 nM in both cases (Fig. 1G, H). These values are in agreement with SPR results showing K_D_ of 4.3 nM (Fig. 1K). In the histone H3 citrullination assay, the addition of heparin allowed full citrullination of substrate already at 200 µM Ca^2+^, whereas it required 2 mM to reach a similar level of modification in the absence of GAG (Fig. 1I). Next, we tested the impact of heparin on PAD4 thermal stability with nanoDSF and observed a stabilizing effect of heparin, which changed the PAD4 melting curve from two distinct unfolding events at 44 and 55°C for PAD4 alone to a single unfolding at 55°C in the presence of heparin at the concentration between 0.5 – 1 µM (Fig. 1J). This observation is consistent with the double denaturation event for PAD4 alone, most likely indicating the initial disruption of multimeric forms at 44°C, which are efficiently stabilized in the presence of heparin, followed by protein unfolding at 55°C.

**Fig. 1.**
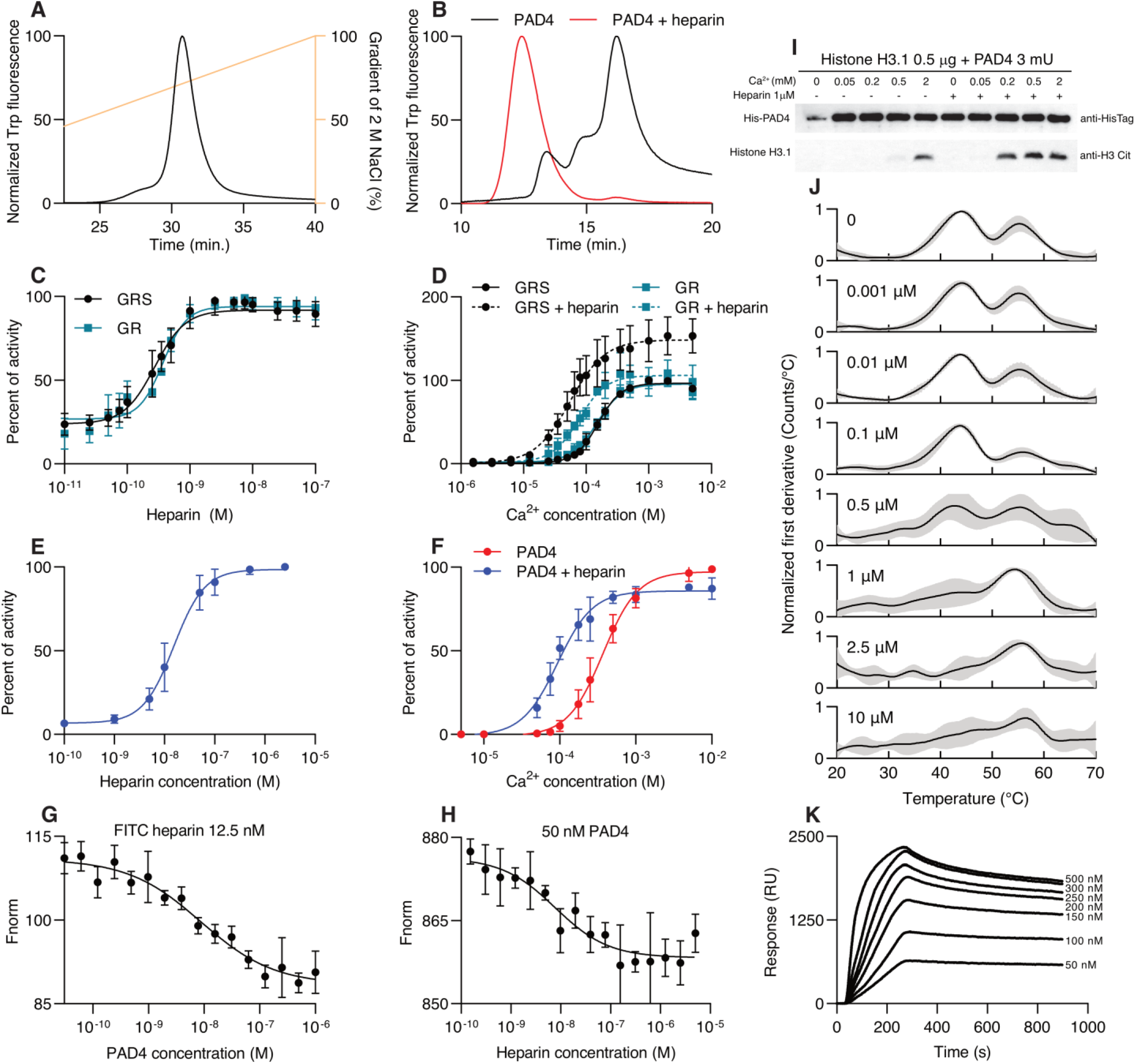
Binding and activation of PAD4 by heparin. (**A**) Elution profile of PAD4 WT from a Poros heparin column in a salt gradient monitored by intrinsic tryptophan fluorescence (λ_ex_ 280 nm, λ_em_ 350 nm). (**B**) Size exclusion chromatography of PAD4 on a Superdex 200 column monitored by tryptophan fluorescence (λ_ex_ 280 nm, λ_em_ 350 nm), PAD4 alone elutes in three forms: monomeric, dimeric and tetrameric (or larger order oligomer), from eluting last to first. After the addition of 5 µM heparin, PAD4 elutes in a high molecular weight complex. (**C** and **D**) Activity of PAD4 measured on peptide substrates Gly-Arg or Gly-Arg-Ser with dansyl fluorescent label, with citrullination detected by HPLC separation of reaction products. (**C**) PAD4 activity in sub-activatory Ca^2+^ concentration (0.1 mM) with the addition of increasing heparin concentrations. Hill equation was fitted to obtain apparent K_D_ values 0.27 ± 0.02 nM and 0.37 ± 0.03 nM for GRS and GR substrates, respectively. Data are presented as percent of maximum activity in the presence of heparin, mean ± SD. (**D**) Activity of PAD4 with/without addition of an activatory heparin concentration (1 µM) and increasing Ca^2+^ concentrations. Data are presented as percent of PAD4 activity in the presence of 2 mM Ca^2+^ as mean ± SD. Hill equation was fitted to calculate Ca_0.5_ values for each substrate and heparin conditions: GRS 155.8 ± 4.2 µM, GRS + heparin 55.1 ± 4.6 µM, GR 148.2 ± 6.4 µM, GR + heparin 77.4 ± 5.3 µM. (**E** and **F**) Activity of PAD4 measured on BAEE substrate and colorimetric detection of citrulline. (**E**) Activation of PAD4 by increasing concentrations of heparin in 0.1 mM Ca^2+^ shown as percent of maximum activity in the presence of heparin. Data are presented as mean ± SD, with Hill equation fitted to obtain apparent K_D_ 15.3 ± 1.4 nM. (**F**) Activation of PAD4 by Ca^2+^ with and without 1 µM heparin, shown as percent of maximum activity in the presence of calcium. Data are presented as mean ± SD with Hill equation fitted to obtain Ca_0.5_ values: no heparin 367 ± 13 µM, with heparin 92.9 ± 4.8 µM. (**G** and **H**) Microscale thermophoresis binding curves of PAD4 and heparin. (**G**) Fluorescently FITC-labelled heparin was at a constant concentration of 12.5 nM and PAD4 was titrated, in (**H**) PAD4 was kept at 50 nM and heparin was titrated. Data are presented as mean ± SD. Hill equation was fitted and gave K_D_ values of 8.8 ± 2.9 nM and 7.7 ± 4.5 nM for (**G**) and (**H**), respectively. (**I**) Western blot assay of PAD4 citrullination of human histone H3.1 at increasing Ca^2+^ concentrations with or without 1 µM heparin. Blot against HisTag of PAD4 is presented as control; results from a representative experiment are shown. (**J**) Thermal unfolding curves of PAD4 with increasing heparin concentrations measured by nanoDSF. Data are shown as mean ± SD. (**K**) Sensograms from a representative SPR interaction assay of various PAD4 concentrations with heparin immobilized on the chip surface. Binding was analyzed with one-site model with estimated K_D_ 4.9 ± 1.1 nM.

### Effect of heparin depends on chain length and charge

After determining the activatory effect of heparin on PAD4, we aimed to investigate which properties of the GAG polymer contribute to this phenomenon. We employed short heparin chain fragments of defined length to verify the size of the molecule required to effectively activate PAD4. In colorimetric assay with BAEE substrate, we determined that PAD4 affinity to heparin decreases with the shortening of the heparin chain (from 16.27 nM for heparin to around 300 nM for Dp12 and even higher values for shorter fragments), in concordance with maximum level of activation (Fig. 2A, C). Similarly, the increase in calcium affinity was less prominent in the presence of smaller heparin fragments (Ca_0.5_ with heparin 92.9 µM and 394.6 µM with Dp4, comparable to 367 µM observed without heparin) (Fig. 2B, C). By plotting the maximum PAD4 activation and Ca_0.5_ values versus oligomer length, we determined that around 12 sugar subunits in the GAG chain are necessary to achieve half of the effect of native heparin (Fig. 2D, E). Furthermore, we used N-desulphated variant of 12 subunits heparin oligomer (Dp12_ND_), which was much weaker activator of PAD4 compared to the unmodified oligomer (Dp12) with full negative charge (apparent K_D_ 741.5 nM for Dp12_ND_ vs 133.2 nM for Dp12 and Ca_0.5_ 259.3 µM for Dp12_ND_ vs 155.8 µM for Dp12) (Fig. 2F, G).

**Fig. 2.**
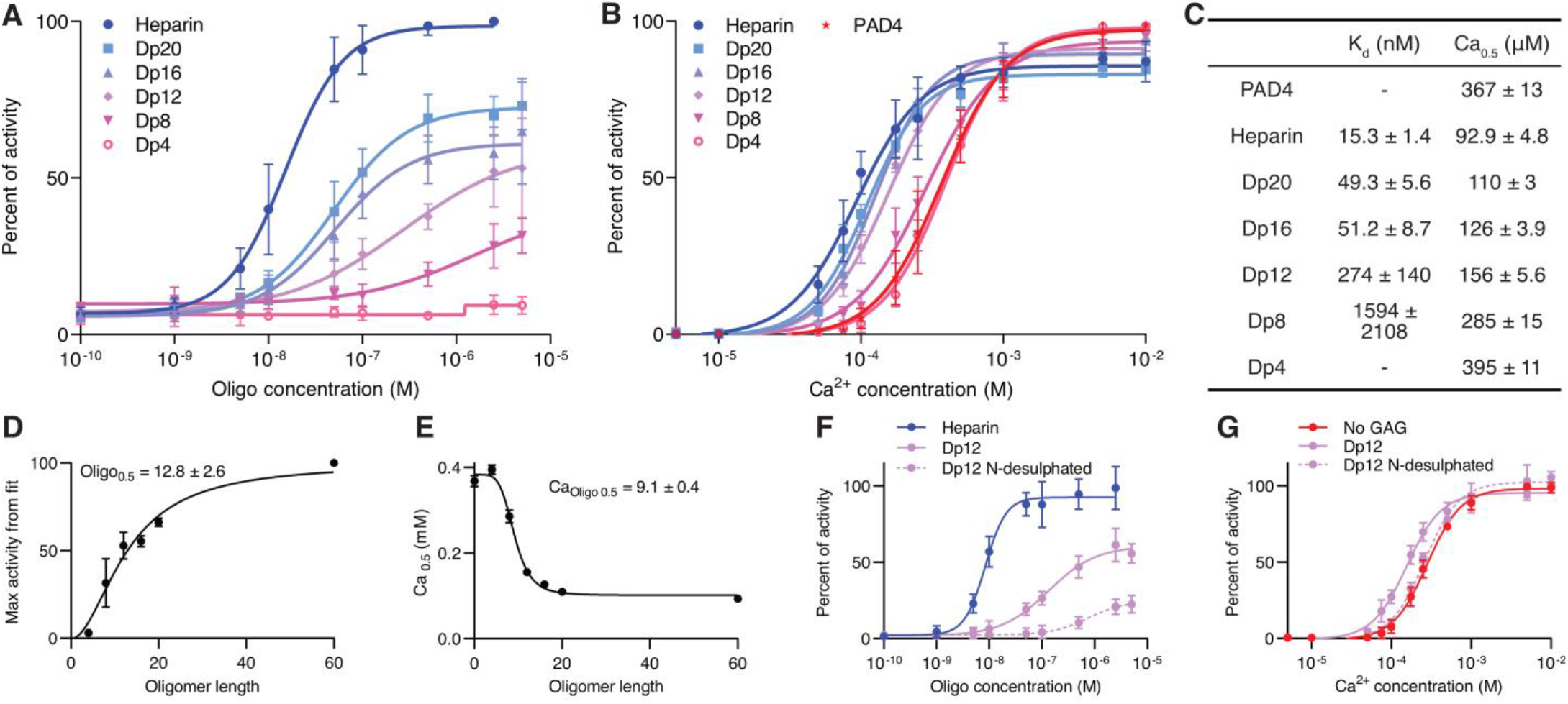
Effect of chain length and charge of heparin oligomers on the activation of PAD4. (**A** and **B**) Activity of PAD4 measured on BAEE substrate with colorimetric detection of produced citrulline. (**A**) Activity of PAD4 in the presence of 0.1 mM Ca^2+^ and heparin oligomers with decreasing length. Data are shown as mean ± SD. Hill equation was fitted to obtain apparent K_D_ values summarized in (**C**). (**B**) Activation of PAD4 by calcium in the presence of 1 µM heparin oligomers of different lengths. Data are shown as mean ± SD and Ca_0.5_ values from Hill equation are shown in (**C**). (**C**) Summary of apparent K_D_ and Ca_0.5_ values for heparin oligomers of various lengths from activity assays in (A) and (B). (**D** and **E**) Theoretical maximum activity of PAD4 from fits in (A) and Ca_0.5_ from (B) in the presence of heparin oligomers vs oligomer length. Hill equation was fitted to determine the length of oligomer at which half of effect is achieved. Error bars represent SE of fitted values. (**F** and **G**) Activity of PAD4 in the presence of heparin oligomer with 12 subunits (Dp12), the same oligomer N-desulphated reN-acetylated (Dp12 N-desulphated), and heparin for comparison, measured with a colorimetric assay and BAEE as substrate, data is shown as mean ± SD. (**F**) Activity in 0.1 mM Ca^2+^ and increasing concentrations of GAGs, Hill equation was fitted to obtain apparent K_D_ values of 8.4 ± 0.6, 133 ± 23 and 741 ± 318 nM for heparin, Dp12, and Dp12 N-desulphated respectively. (**G**) Activation of PAD4 with Ca^2+^ in the presence of 1 µM of GAGs, Ca_0.5_ values from fitted Hill equation: 280.8 ± 7.5, 155.8 ± 5.3, 259.3 ± 9.5 µM for no GAG, Dp12 and Dp12 N-desulphated respectively.

### Mechanism of PAD4 activation by heparin

Based on the structural data of charge distribution on the surface of the protein we aimed to identify the sites in PAD4 that are likely responsible for binding and/or activation of the enzyme by heparin. We have shown the importance of the negative charge of heparin for efficient activation of PAD4, therefore we narrowed the search for the binding site by mutagenesis only to positively charged regions. We prepared two mutants of the regions suspected to be involved in binding of heparin oligomers (HLS: K126S K128S R131S K134S R137S and NLS: K59S K60S K61S K81S K91S) and tested their binding to an analytical heparin column. Both mutants eluted earlier in gradient compared to PAD4 WT, suggesting weaker binding (Fig. 3A). Similarly, when analyzed by size exclusion chromatography, both mutants showed only subtle changes in elution profiles after the addition of heparin, with limited formation of larger oligomers in the HLS mutant. Interestingly, both mutants even in the absence of heparin eluted predominantly as monomeric and possibly tetrameric forms even in the absence of heparin, and no dimerization was observed (Fig. 3B). Both mutants showed weaker activation by heparin in low calcium concentrations, however apparent K_D_ values were similar compared to WT (HLS 7.5 nM, NLS 5.7 nM, WT 6.9 nM) (Fig. 3C, D). These values were confirmed in SPR analysis, which showed a marginally weaker binding of mutants to immobilized heparin compared to the WT enzyme (Fig. 3C). These results indicate that the identified sites have only a limited effect on the PAD4-heparin interaction when analyzed separately. Activity of PAD4 was reported to be dependent on the dimerization of the protein. Therefore, we prepared mutants of PAD4 that were previously reported to have significantly impaired dimerization ability: R8E, Y435A, and double mutant to test activation of monomeric PAD4 by heparin (*10*–*12*). All three dimerization mutants eluted earlier than WT or HLS, NLS mutants from heparin column, suggesting even weaker interaction (Fig. 3A). Size exclusion analysis showed the three mutants are almost exclusively present as monomers with a minor dimeric fraction. The addition of heparin resulted in the detection of only a small amount of larger complexes (Fig. 3B). Further, in the activation assay, mutants R8E and Y435A show very weak activation, and the double R8E Y435A mutant was not activated by heparin, indicating dimerization as the essential feature of the heparin binding. Obtained apparent K_D_ values were still in low nanomolar range (13 nM vs 7 nM for WT) however, due to the low levels of activity, they were determined with low confidence. When tested with SPR, all three mutants were able to bind to the heparin-coated chip, but the measured K_D_ values were higher than WT (Fig. 3D), and the signal response was low, likely indicating binding of only the residual dimer fraction.

**Fig. 3.**
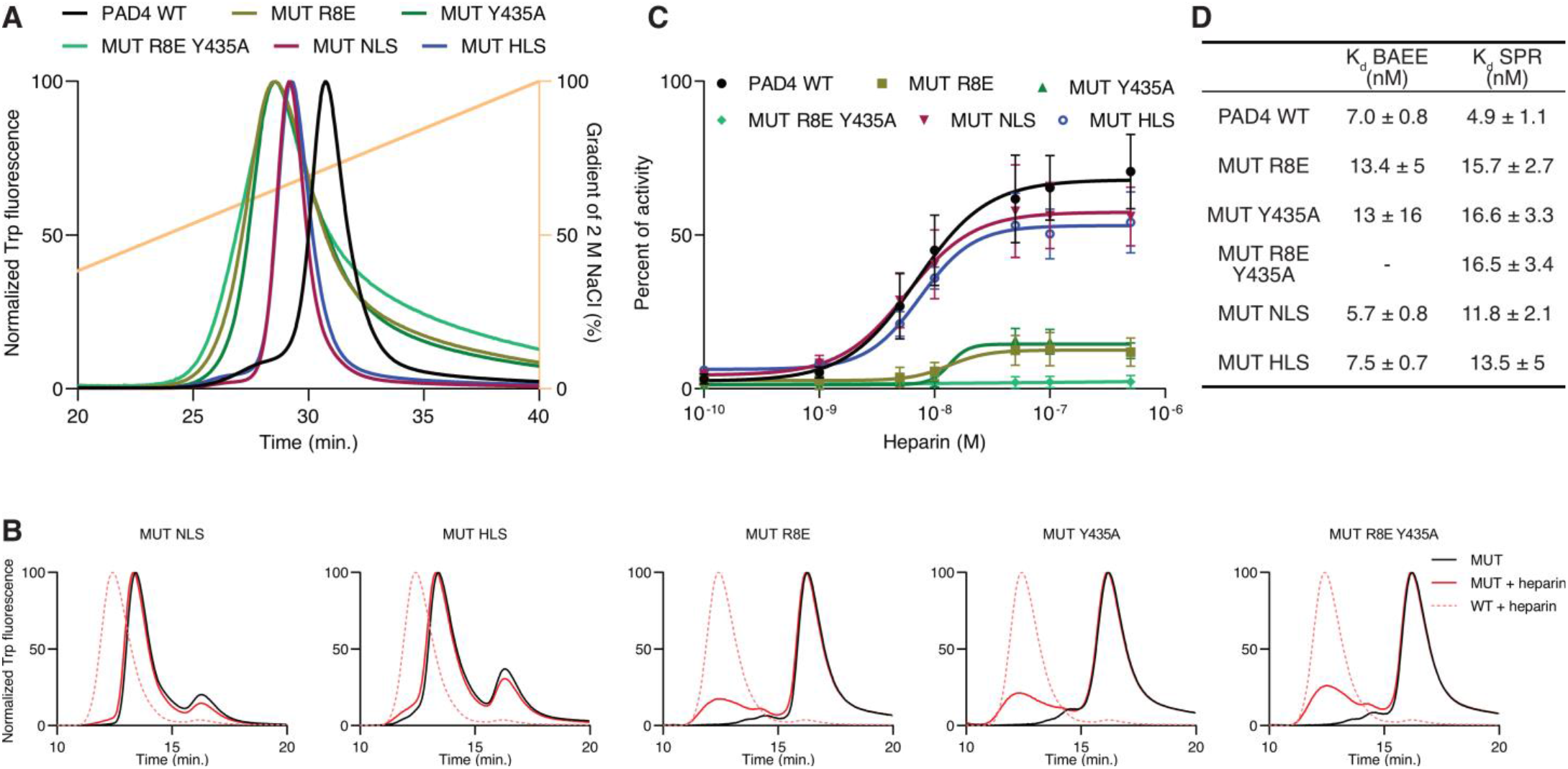
Characterization of heparin binding to PAD4 mutants of dimerization (R8E, Y435A, R8E Y435A) and regions involved in interaction with heparin (HLS: K126S K128S R131S K134S R137S and NLS: K59S K60S K61S K81S K91S). (**A**) Elution profiles of PAD4 WT and mutants from a Poros heparin column in a salt gradient monitored by intrinsic tryptophan fluorescence (λ_ex_ 280 nm, λ_em_ 350 nm). (**B**) Size exclusion chromatography of PAD4 mutants with and without 5 µM heparin on a Superdex 200 column monitored by tryptophan fluorescence (λ_ex_ 280 nm, λ_em_ 350 nm). PAD4 WT with heparin is shown for reference. (**C**) Activation of PAD4 and mutants with heparin in 0.1 mM of Ca^2+^ with BAEE substrate and colorimetric citrulline detection. Activity is presented as percent of activity of each mutant in 5 mM Ca^2+^, data are presented as mean ± SD. Hill equation was fitted to calculate apparent K_D_ for each mutant showed in (**D**) together with K_D_ from SPR assay of interaction with immobilized heparin.

### Biological properties of PAD4 activation with GAG

We investigated whether natural GAGs other than heparin have an effect on PAD4 activation. We found that among dermatan sulphate, heparan sulphate, and chondroitin sulphates A, C, D all tested compounds activated PAD4 with varying potency at 1 µM concentration in presence of 0.1 mM Ca^2+^ (Fig. 4A). Dermatan and heparan sulphates activated PAD4 to the same level as heparin, while chondroitin sulphates induced lower activity. All compounds stimulated citrullination of histone H3 at the same concentrations of GAG and Ca^2+^ as in the presence of heparin, while no modification was present under the same conditions without GAG (Fig. 4B). Further, we tested binding of PAD4 to naturally occurring GAGs on the surface of CHO cells using FACS analysis and mutant cell lines with impaired GAG production (*24*–*26*) (Fig. 4C). We found that both GAG-deficient cell lines (650 and 745) bound PAD4 less efficiently. This observation confirms that ability to bind to PAD4 is not restricted to free GAG chains but can also occur on the surface of cells. The effect of heparin on neutrophils was tested using isolated human PMN cells (hPMN) with DNA release assay and staining of citrullinated histone H3, both being constituents of NETs. Heparin induced concentration-depended release of DNA over the course of 4 hours and stimulated citrullination of histone H3 in hPMNs (Fig. 4D and E). Both of those effects can be attributed to the activation of PAD4.

**Fig. 4.**
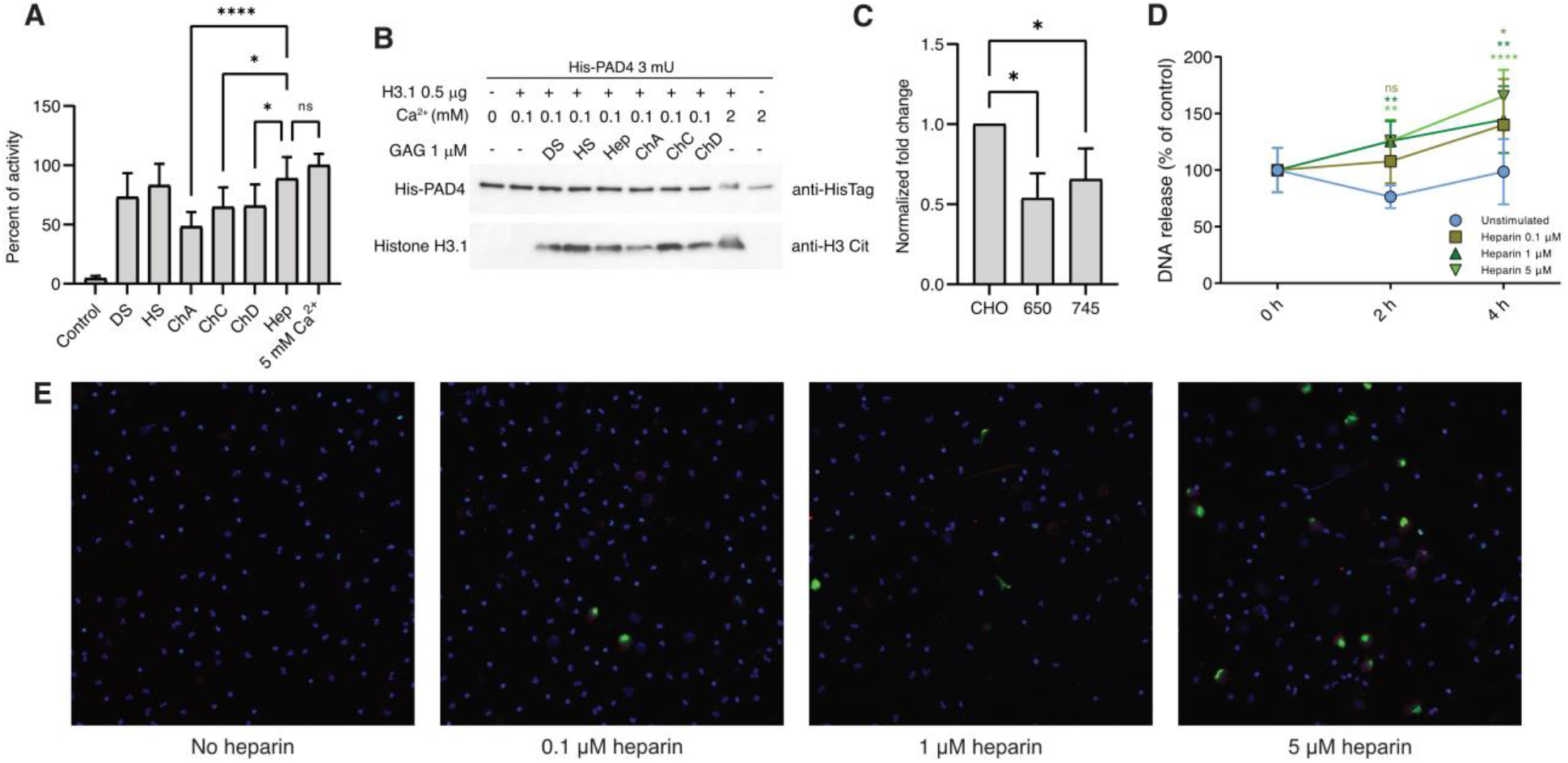
Biological properties of PAD4 activation with GAG. (**A** and **B**) Activation of PAD4 in 0.1 mM Ca^2+^ after the addition of 1 µM of GAG (DS-dermatan sulphate, HS – heparan sulphate, ChA – chondroitin A, ChC – chondroitin C, ChD – chondroitin D, Hep – heparin). (**A**) Colorimetric PAD4 activity assay with BAEE substrate. Data is presented as mean ± SD as percent of activity in 5 mM Ca^2+^. Statistical significance was tested with one-way ANOVA with Tukey multiple comparison correction, ns – not significant, ^*^ - p < 0.05, ^****^ - p < 0.0001. (**B**) Western blot assay of PAD4 citrullination of human histone H3.1. Blot against HisTag of PAD4 is presented as a control; results from representative experiment are shown. (**C**) FACS analysis of PAD4 binding to CHO-K1 wild type cells and its mutated cell lines with impaired GAG production (PgsB-650 and PgsA-745). Results are presented as signal fold change over background of cells not treated with PAD4 and normalized to control CHO cells. Data is presented as mean ± SD, statistical significance was tested with one-way ANOVA with Dunnett multiple comparison correction, ^*^ - p < 0.05. (**D**) Release of DNA from hPMN cells stimulated with increasing concentrations of heparin. Data is presented as mean ± SD, of signal normalized to time 0. Statistical significance was tested with two-way ANOVA with Tukey multiple comparison correction, at each time point significance compared to unstimulated cells is shown, ns – not significant, ^**^ - p < 0.01, ^****^ - p < 0.0001. (**E**) Immunocytochemistry staining of hPMNs stimulated with heparin for 4 hours at indicated concentrations. Blue – DNA, green – citrullinated histone H3, red – PAD4.

## Discussion

The involvement of PAD4 in the pathogenesis of RA is a well-established concept in the study of this disease (*7, 19, 20*). The presence of antibodies against citrullinated peptides is an important prognostic and diagnostic tool in RA, and the activity of PAD4 is a prime candidate for the source of such peptides. This hypothesis is supported by reports indicating presence of both, active PAD enzymes and citrullinated epitopes directly in the synovial fluid of patients (*27, 28*). This is often attributed to PAD4 due to its presence in NETs, which can be released under the inflammatory state (*2, 3, 7, 20*). Biochemical analysis of PAD4 showed that activation of this enzyme relies on calcium concentration, but blood levels of calcium seem too low to achieve maximum activation (*1, 13, 16, 17*). This observation prompts the search for other modulators of PAD enzymes, and PAD4 activity in particular. Based on the reports that heparin and other GAGs are also able to cause RA-like symptoms and that heparin induces neutrophils to release NETs, contributing to heparin-induced thrombocytopenia, we hypothesized that these factors may be linked by activation of PAD4 (*21*–*23*). Indeed, we showed that heparin activates PAD4 with low nanomolar affinity. The strength of this interaction seems to be the result of very slow complex dissociation, as demonstrated by SPR analysis, and PAD4 could only be released from heparin resin under high salt concentrations. Activation of PAD4 by heparin was achieved through the increase of the PAD4’s affinity for calcium, allowing full activation of PAD4 well below 1 mM Ca^2+^. Such levels can be found in physiological fluids, and nanomolar concentrations of heparin would then trigger PAD4 to activate, forming a stable complex that could remain active over time. A distinctive feature of heparin is its polymeric structure and strong negative charge; in our study both of those factors are essential for the activation of PAD4. Short or undersulphated fragments had a weaker effect on the enzyme, similar to observations made for other GAGs interacting with proteins (*29*). Further study of PAD4 binding to full-length heparin revealed formation of high molecular weight and high hydrodynamic radius complexes in size exclusion chromatography, consistent with heparin-bound tetramer. Mutagenesis study of suspected binding sites of PAD4 was not fully conclusive, as mutants of two putative binding sites were still activated by heparin to some extent; however, PAD4 dimer formation and the presence of heparin complexes were limited in SEC for these variants. We speculate that heparin binding occurs on positively charged surfaces of PAD4, widely distributed throughout the structure, and the removal of only some of them by mutagenesis has a limited effect. Furthermore, analysis of PAD4 mutants with impaired dimerization showed a much more significant reduction of enzyme activation by heparin. Based on that, we propose that heparin binds to already dimerized PAD4, resulting in the formation of stable large complexes and lowers the calcium activation threshold through allosteric regulation, potentially involving multiple dimers connected by heparin chains. A similar mechanism of activation was recently described using selected monoclonal antibodies that stabilized PAD4 dimer and increased its activity, while blocking dimerization lowered activity (*12*). Earlier studies revealed that a fraction of RA patients possess the cross-reactive anti-PAD3/PAD4 antibodies, which increase enzyme activity by increasing calcium affinity, and their presence correlates with the disease progress (*30*– *32*). This mode of PAD4 activation was also investigated recently by generation of synthetic peptides with similar effect (*33*). Herein, we identify GAGs as the first ever physiological activators of PAD4. The anti-PAD3/4 antibodies mentioned earlier are present in only a minor fraction of RA patients. The universal presence of GAGs in all individuals might therefore contribute to the explanation of RA development in a larger number of patients. The effect of GAGs on neutrophil biology is further supported by the increased DNA release upon heparin stimulation, correlated with the increase in histone H3 citrullination, which are markers of NETosis. Similar observations were made previously, as heparin was found to induce vital NETosis in human neutrophils, and NETosis itself is the main driver of thrombosis in heparin-induced thrombocytopenia (*23, 34*). Reported herein heparin-induced PAD4 activation resulting in downstream NETosis could partially explain the mechanism underlying life-threatening HIT. The biological importance of PAD4 activation is highlighted by the observation that this property is not unique to heparin, but other GAG molecules have similar properties. This universal character of GAG involvement in PAD4 binding was demonstrated by our CHO-cell analysis, as GAG-deficient cell lines were incapable of efficient PAD4 binding, as demonstrated by flow cytometry. Importantly for the RA pathogenesis, chondroitin sulphates were efficient activators of PAD4. We hypothesize that cartilage-derived GAG-induced overactivation of PAD4 might lead to or exacerbate the production of citrullinated peptides in the synovial fluid following the release of PAD4 from neutrophils during NETosis. This might be an early step leading to RA development after initial inflammatory event in the joint or other site in the body, as well as a driving force of joint destruction providing citrullinated peptides sustaining the immune response in synovial fluid. In conclusion, we provide the first evidence of naturally occurring coactivators of PAD4 other than calcium, which can help explain RA pathogenesis as well as the effects of heparin on neutrophils.

## Funding

Research was supported by Norway Financial Mechanism for years 2014 - 2021 under the GRIEG Project (2019/34/H/NZ1/00674) to Tomasz Kantyka, PhD.

National Science Centre, Poland, Harmonia 10 (2018/30/M/NZ1/00367) to Ewa Bielecka, PhD.

Research Council of Norway and National Institutes of Health (296129) to Piotr Mydel, PhD.

## Author contributions

Conceptualization: TK, EB, GB

Data curation: PW, AB, TK

Methodology: GPB, TK, EB, AB, ŁP, JN, JK

Investigation: GPB, EB, KM, ŁP, AB, PW, MK, DK, JN

Visualization: GPB, EB, ŁP, JN, EW, MK, AB, PW

Funding acquisition: TK, PM, MP, EB

Project administration: TK, PM, MP, PG

Supervision: PG, JK, PM, MP, TK

Writing – original draft: GPB

Writing – review & editing: GPB, EB, ŁP, TK, PG, PM, MP

## Supplementary Materials

Materials and Methods

