## Supplementary Materials for "Glycosaminoglycans act as activators of peptidylarginine deiminase 4"

**Materials and Methods**

Recombinant PAD4 expression and purification

Full length human PAD4 sequence with N-terminal affinity Histidine tag: MGHHHHHHHHHHH cloned into pET16b vector was synthesized by GenScript Biotech Corporation. Subsequent mutant variants of PAD4 were also synthesized by GenScript in the same expression vector: HLS (K126S K128S R131S K134S R137S); NLS (K59S K60S K61S K81S K91S); R8E; Y435A; R8E Y435A. Received plasmids (10 ng) were transformed into *E. coli* BL21(DE3) expression strain with heat shock protocol, plated on LB agar plates with 100 µg/ml ampicillin and incubated at 37°C. For expression three single colonies were inoculated into 50 ml of LB with 100 µg/ml ampicillin and incubated overnight at 37°C with shaking 180 rpm. The next day, culture was diluted 50x in 800 ml of fresh LB with 100 µg/ml ampicillin (expression was scaled up when necessary, typically 2 cultures of 800 ml were used for one purification batch) and incubated at 37°C with 180 rpm shaking until OD_600_ reached 0.6. Cultures were then incubated for 20 minutes at 4°C, expression was induced with 0.5 mM IPTG (isopropyl β-D-thiogalactopyranoside, BioShop #IPT001) and carried for 16 hours at 26°C with 180 rpm shaking. Cultures were centrifuged at 6500 g, 4°C, for 20 minutes and pelleted bacteria were collected and stored at -20°C until further purification. Cells were suspended in lysis buffer: 50 mM sodium phosphate buffer pH 7.5, 0.5 M NaCl, 0.5 mM EDTA, 10% glycerol, 0.1% Triton X-100, 10 mM imidazole on ice and sonicated for 3 minutes total pulse with pulse cycle 5 seconds on, 5 seconds off with 70% amplitude on Sonics VCX500 sonicator with 13 mm tip. Lysate was then centrifuged at 40000 g, 4°C, 40 minutes and clarified supernatant was loaded on 5 ml HisTrap FF column (Cytiva #17531901) connected to Akta Pure system (Cytiva). Column was then washed with 5 column volumes of washing buffer (50 mM sodium phosphate buffer pH 7.5, 0.5 M NaCl, 0.5 mM EDTA, 10% glycerol, 10 mM imidazole) and protein was eluted with linear gradient 0 to 100% of 500 mM imidazole in washing buffer over 6 column volumes. Fractions containing PAD4 were then subjected to buffer exchange to binding buffer: 50 mM Tris pH 7.5, 300 mM NaCl, 5% glycerol, 0.5 mM EDTA on HiPrep 26/10 Desalting column (Cytiva #17508701). PAD4 was then loaded on HiTrap Heparin HP column (Cytiva #17040601), washed with 10 column volumes of binding buffer and eluted with linear gradient 0 to 100% of 2 M NaCl in binding buffer over 15 column volumes. Protein containing fractions were pooled and subjected to size exclusion chromatography on HiLoad 16/600 Superdex 200 pg column (Cytiva #28989335) in 50 mM Tris pH 7.5, 300 mM NaCl. Finally, protein was concentrated on Amicon Ultra 15, 10000 NMWL (Millipore #UFC901024) to 3 - 5 mg/ml and stored at -80°C.

Affinity chromatography of PAD4 and its mutants on heparin column

Binding of PAD4 and its mutants to heparin column was performed on a POROS Heparin 50 μm 2.1 x 30 mm column (Thermo Scientific #4333411) connected to a Shimadzu Nexera 2 chromatography system. PADs at 0.25 µM concentration were applied on column in buffer: 50 mM Tris pH 7.5, 150 mM NaCl and resolved in 7.5-100% gradient of phase B (50 mM Tris, pH 7.5, 2 M NaCl) over 30 column volumes. Tryptophan fluorescence of samples was monitored at excitation wavelength of 280 nm and emission wavelength of 350 nm.

SEC analysis of PAD4 and its mutants in complexes

Analytical SEC analysis of PAD4 and its mutants in the presence or absence of 5 µM heparin (Heparin sodium salt from porcine intestinal mucosa, Sigma-Aldrich #H3149) were performed on a Superdex 200 Increase 3.2/300 column (Cytiva, #28990946) connected to a Shimadzu Nexera 2 chromatography system. Each protein (30 µl) was run at 0.15 µM concentration in running buffer consisting of 50 mM Tris pH 7.5, 300 mM NaCl, 0.1 mM CaCl_2_, 0.05% Tween20, 0.25 mM DTT. Tryptophan fluorescence of samples was monitored at excitation and emission wavelength of 280 and of 350 nm respectively. Retention time of each peak was compared with protein standards from the HMW Gel Filtration Calibration Kit (Cytiva #28403842).

RP-HPLC based method for measurement of PAD4 activity in the presence of different calcium and heparin concentrations

PAD4 activity in the presence of different CaCl_2_ or heparin concentration was measured with RP-HPLC based method. PAD4 (0.25 mU) was incubated with CaCl_2_ concentrations 0-5 mM (calcium curve) in presence or absence of 0.025 µM heparin (Heparin sodium salt from porcine intestinal mucosa, Sigma-Aldrich #H3149); or heparin concentrations 0-2.5 µM (heparin curve) in the presence of 0.1 mM CaCl_2_. Fluorescently labelled substrate was used at 100 µM: Dansyl-Gly-Arg or Dansyl-Gly-Arg-Ser (custom synthesized by PeptideWeb) in reaction buffer (100 mM Tris pH 7.5, 5 mM DTT, 1% DMSO), samples were incubated for 1 hour at 37°C in 50 µl total volume. Reaction was stopped by the addition of 5 µl 1% TFA. Samples were resolved on an Aeris 2.6 µm Peptide XB-C18 50 x 2.1 mm column (Phenomenex #00B-4505-AN) connected to a Nexera 2 chromatography system (Shimadzu) in gradient of phase A (0.1% TFA in water) and B (80% acetonitrile, 0.08% TFA in water) for Dansyl-GR 5-20%, for Dansyl-GRS 5-30% of phase B over 26 column volumes. Dansyl-label fluorescence was monitored at excitation wavelength of 333 nm and emission wavelength of 533 nm. Area of the peaks corresponding to the citrullinated products were calculated with LabSolution Software (Shimadzu). Results are shown as a percent of PAD4 activity on each substrate in 2 mM CaCl_2_ without heparin for calcium curve and as percent of maximal PAD4 activity on each substrate in the presence of heparin for heparin curve.

PAD4 activity assay with BAEE substrate

PAD4 activity was measured with spectrophotometric COLDER Assay (*35*). For general evaluation of PAD4 activity, enzyme was incubated in 50 µl reaction volume in buffer containing 100 mM Tris pH 7.5, 5 mM DTT, 5 mM CaCl_2_, 2% DMSO and 10 mM Nα-Benzoyl-L-arginine ethyl ester hydrochloride substrate (BAEE, Sigma-Aldrich #B4500) for 1 hour at 37°C. The reaction was stopped by the addition of 10 µl 5 M HClO_4_. For the development of color reaction product, 150 µl of developing solution (containing 0.16% diacetyl monoxime, 0.0033% thiosemicarbazide and 0.16 mg/ml FeCl3 (all from Sigma-Aldrich) in 16.33% H_2_SO_4_ and 11.33% H_3_PO_4_ (Avantor Performance Materials Poland)) was added to the samples and incubated at 110°C for 3 minutes. Absorbance of the samples was measured at 535 nm with a Hidex Sense multiplate reader (Hidex), and compared to the standard curve of L-citrulline (Sigma-Aldrich #C7629), 1 mU of PAD4 activity is defined as 1 nanomole of citrulline produced by the enzyme within 1 hour of incubation at 37°C.

Evaluation of PAD4 and PAD4 mutants activity in the presence of different concentrations of heparin/GAGs/Dp forms (GAG curves) was performed in similar manner. Briefly, ~30 mU of PAD4 were incubated (37°C, 1 hour) in activity buffer described above but with 0.1 mM CaCl_2_ with addition of increasing heparin/GAGs/Dp concentrations 0-5 µM. Later on the reaction was stopped and developed as above. Activity of PAD4 in the presence of different activator concentrations was calculated in accordance to L-citrulline standard curve and presented as percent of maximal PAD4 activity in the presence of heparin.

Evaluation of PAD4 activity in the presence of different calcium concentrations (Calcium curve) with and without heparin/Dp was performed analogically. Briefly, ~30 mU of PAD4 was incubated in reaction buffer containing 0-10 mM CaCl_2_ alone or with 1 µM Heparin/Dp for 1 hour at 37°C. Later on, the reaction was stopped and developed as above. Activity of PAD4 in the presence of different calcium concentrations was calculated in accordance to L-citrulline standard curve and presented as percent of maximal PAD4 activity in calcium alone.

Materials used: Heparin sodium salt from porcine intestinal mucosa, Sigma-Aldrich #H3149; GAGs and heparin oligomers all from Iduron: Dp4 (#HO04), Dp8 (#HO08), Dp12 (#HO12), Dp12 N-desulphated reN-acetylated (#reAc DSHO12), Dp16 (#HO16), Dp20 (#HO20), dermatan sulphate (#GAG-DS01), heparan sulphate (#GAG-HS01), chondroitin sulphate A (#GAG-CSA01), chondroitin sulphate C (#GAG-CSC01), chondroitin sulphate D (#GAG-CSD01).

Histone citrullination

For each reaction, 0.5 µg of human histone H3.1 (Sigma-Aldrich #H2292) was incubated for 2 hours at 37°C with 3 mU of PAD4 in the presence or absence of 1 µM GAG (Heparin sodium salt from porcine intestinal mucosa, Sigma-Aldrich #H3149) in 50 mM Tris pH 7.5, 5 mM DTT and 0-2 mM CaCl_2_. The reaction was stopped by addition of 6x concentrated sample buffer and boiled for 15 min. at 100°C. Samples were then subsequently analyzed by western blot. Briefly, after SDS-PAGE on 10% polyacrylamide gels, proteins were transferred on PVDF membrane (Cytiva #10600023). Membranes were blocked with 5% skim milk (BioShop #SKI400) in washing buffer 50 mM Tris pH 7.5, 150 mM NaCl, 0.05% Tween20 for 3 hours at RT. Then membranes were cut based on molecular weight marker and the bottom part of the membrane (containing histone H3.1) was stained with primary rabbit antibodies anti-histone H3 Cit (Abcam #ab5103, dilution 1:2000) overnight at 4°C, followed by staining with secondary antibodies goat anti-rabbit-HRP (Sigma-Aldrich #A0545, dilution 1:20000) for 2 hours at RT. The upper part of the membrane (containing HisTag-PAD4) was stained with mouse monoclonal anti-polyHistidine−HRP antibody (Sigma-Aldrich #A7058, dilution 1:20000) for 2 hours at RT. HRP signal was detected on X-ray film blue (AGFA #CP-BU NEW) with Pierce ECL substrate (Thermo Scientific #32106). For the evaluation of other GAGs influence on histone citrullination, the procedure was performed analogously but in the presence of 0.1 mM calcium chloride and respective GAG at 1 µM.

Microscale thermophoresis (MST)

Microscale thermophoresis assays were done in 20 mM Tris pH 7.5, 300 mM NaCl, 0.05% Tween20. In the first setting, the signal was measured from FITC-Heparin (Invitrogen, Heparin Fluorescein Conjugate #H7482) at a final concentration of 12.5 nM and PAD4 was added at final concentrations starting at 1 µM followed by 15 twofold dilutions. Samples were mixed, centrifuged at 16000 g, 4°C, 5 minutes and loaded into Monolith Premium Capillaries (NanoTemper #MO-K025). Samples were measured on a Monolith NT.115 (NanoTemper) instrument using nano-blue laser at 100% power. Data from initial fluorescence were used for further analysis. In the second setting, concentration of PAD4 was kept constant at 50 nM and heparin (Heparin sodium salt from porcine intestinal mucosa, Sigma-Aldrich #H3149) was added at final concentrations starting at 5 µM followed by 15 twofold dilutions. Samples were mixed and centrifuged as before, and loaded into Monolith LabelFree capillaries (NanoTemper #MO-Z022). Measurements were done on a Monolith NT.LabelFree (NanoTemper) instrument with Labelfree laser at 80% power and medium MST setting and data from MST mode were used for analysis. In both experiments data were imported into MO.Affinity Analysis 2.3 software and exported for further analysis and fitting of Hill slope.

Protein stability tests in the presence of heparin with nanoDSF

PAD4 was used in a final concentration 0.2 mg/ml in assay buffer 50 mM Tris pH 7.5, 150 mM NaCl with varying concentrations of heparin (0 – 10 µM) (Heparin sodium salt from porcine intestinal mucosa, Sigma-Aldrich #H3149). Samples were vortexed and centrifuged at 16000 g, 4°C, 5 minutes and loaded into Prometheus High Sensitivity Capillaries (NanoTemper #PR-C006). Thermal stability was measured on Prometheus Panta (NanoTemper) in range 20 - 95°C with 1°C/minute ramp. Data was analyzed with NanoTemper analysis software, exported and processed using Origin software v2023 (OriginLab). The processing steps included first-order derivatization with Savitzky-Golay (SG) smoothing (polynomial order 2, points window 20), baseline subtraction using the Origin Peak Analyzer with the baseline detection ALSS method (threshold 0.1, smoothing factor 6, iterations 10), normalization of the data, and final smoothing using the SG method (order 2, points window 30).

Heparin-PAD4 interaction analysis with SPR

For immobilization on the SPR chip, heparin was first biotinylated. For biotinylation reaction 2 mg of heparin (Heparin sodium salt from porcine intestinal mucosa, Sigma-Aldrich #H3149) and 2 mg of EZ-Link Amine PEG3-biotin (Thermo Scientific #21347) were dissolved together in 200 µl HPLC grade water. The mixture was then added to 10 mg of NaCNBH_3_ (sodium cyanoborohydride, Sigma-Aldrich #156159) and incubated at 70°C for 24 hours, with an additional 10 mg of NaCNBH_3_ was added and incubated for another 24 hours. Samples were concentrated to 50 µl on Pierce Concentrator PES 3K MWCO (Thermo Scientific #88525), diluted to 500 µl with 50 mM Tris pH 7.5 and concentrated again. Dilution and concentration were repeated 6 times in total to remove leftover reagents. Finally, biotinylated heparin was brought to 200 µl. Interaction analysis by surface plasmon resonance was performed using OpenSPR two channel device (Nicoya) and Biotin-Streptavidin Sensor Kit (Nicoya #SEN-AU-100-10-STRP-KIT). Assay was performed in buffer 100 mM Tris pH 7.4, 200 mM NaCl, 0.2 mM CaCl_2_. The SPR chip was loaded into instrument and prepared according to manufacturer instructions, biotinylated heparin was immobilized by single injection of 500 µg/ml in assay buffer. Dilutions of PAD4 or respective mutants were prepared in assay buffer at indicated final concentrations. Injections were performed with flow 20 µl/min., contact time 300 s, dissociation time 600 s. After each injection of PAD4 chip was regenerated with injection of 2 M NaCl in assay buffer, flow 150 µl/min, contact 40 s, dissociation 270 s. Sensograms were imported and analyzed in TraceDrawer software ver. 1.9.1, data were fitted with 1:1 diffusion corrected model and bulk shift set to constant 0.1.

FACS analysis of PAD4 binding to CHO cells

CHO cell line (CHO-K1, ATCC #CCL-61) and its mutant lines with disrupted glycosaminoglycans synthesis: 650 (pgsB-650, ATCC #CRL-2243) and 745 (pgsA-745, ATCC #CRL-2242) (*24*–*26*) were cultured in DMEM F-12K Nut Mix (with 10% FBS and 0.5% penicillin/streptomycin). CHO cell cultures were detached from the culture flasks using Accutase (Thermo Scientific #00-4555-56) and washed once with FACS buffer (PBS containing 1% BSA). Cells were counted in the Burker chamber after Trypan blue staining, and resuspended in FACS buffer to the concentration of 30 mln/ml. Cells were allowed to rest for 30 minutes prior to staining initiation. 100 µl of cell suspension was mixed with recombinant PAD4 at final concentration of 0.1 mg/ml and incubated for 1 hour at 37°C. Cells were then centrifuged (300 g, RT, 8 min.) and washed with FACS buffer. Bound PAD4 was detected via primary antibodies – mouse anti-PAD4 (Abcam #ab128086, dilution 1:100) for 30 minutes at room temperature. Cells were washed with FACS buffer and further incubated with secondary antibodies – rat anti-mouse FITC-conjugated antibody (BD #562028, dilution 1:200) for 30 minutes at room temperature. Samples were diluted to final volume of 200 µl and analyzed on BD LSRFortessa Cell Analyzer flow cytometer (BD). Fcs files were then processed in FlowJo (BD). First, debris was gated out in FSC-SSC plot. In the next step, single cells were selected via FSC-A – FSC-H analyses. Then, FITC histograms were prepared. Median fluorescence intensity (MFI) of signal from the single-cell gate was used to calculate the rate of PAD4 binding in relation to the “baseline” signal recorded for cells stained only with the secondary antibody. Results are presented as fold FITC signal increase over the baseline MFI. Experiment was repeated three times, each time with two technical replicates per condition.

Human PMN cells experiments

PMN isolation

Neutrophils were isolated from peripheral blood collected from de-identified human healthy donors from the Red Cross (Kraków, Poland), thus the study was excluded from human subject approval. Neutrophils were isolated within 1 hour post bleeding according to the procedure described previously (*36*). Briefly, as a result of the separation of blood components in a density gradient using Pancoll (1.077 g/ml density; Pan Biotech #P04-63500) a fraction rich in neutrophils and erythrocytes was obtained. Then neutrophils were separated after a 30 minute incubation with 1% polyvinyl alcohol - PVA (Avantor Performance Materials Poland #734892424). Neutrophils were collected from the upper layer and after centrifugation (280 g, 10 min at RT) the residual erythrocytes were removed by lysis in water. Purified neutrophils were suspended in serum-free DMEM medium without phenol red (Thermo Scientific #21063029).

Release of DNA

The release of DNA from neutrophils in response to different concentrations of heparin (Iduron #HEP001/100) was determined using 96-well black, clear flat bottom microtiter plates (VWR, #732-3737). All assays were run in three independent biological replicates, each run in three technical replicates. hPMNs (2×10^5^) were incubated in a humidified chamber for 60 minutes at 37°C, 5% CO_2_ to allow cell adhesion. Subsequently, medium (control) or heparin (0 – 5 µM) were gently added to the cells, and cells were further incubated for 2 and 4 hours. After cell stimulation 5 μM SYTOX Orange (Invitrogen #S11368) was added, staining was allowed to proceed for 5 min, and fluorescence in each well was determined as arbitrary fluorescence units (AFU, excitation 547 nm, emission 570 nm) using Hidex Sense multiplate reader (Hidex). DNA release was also measured in neutrophils just after their isolation (time 0).

Immunocytochemical staining

hPMNs (1.5×10^5^) were incubated in a humidified chamber on circular glass slides that were placed at the bottom of the wells in a 24-well plate for 60 minutes at 37°C, 5% CO_2_ to allow for cell adhesion. Subsequently, medium (control) or heparin (0 – 5 µM) (Iduron #HEP001/100) were gently added to the cells, and cells were further incubated for 4 hours. Immediately after incubation, cells were carefully fixed sequentially in 1%, 2% and 3% paraformaldehyde in PBS for respectively 2, 10 and 20 minutes and washed in PBS to not disrupt formed NETs. Slides were then incubated in blocking solution containing 3% BSA (bovine serum albumin, Sigma-Aldrich #A9647) in PBS. Slides were then labelled with the rabbit primary poly-clonal anti-citrullinated histone H3 antibody (Abcam #ab5103 dilution 1:400) and the mouse anti-PADI4 clone OTI4H5 antibody (Origene #CF504813 dilution 1:1000) and incubated overnight at 4°C. The slides were subsequently washed in PBS and incubated with the secondary antibodies: Alexa Fluor 488 chicken anti-rabbit IgG (H+L) (Invitrogen #A21441 dilution 1:500) and Alexa Fluor 647 goat pAb to Ms IgG (Abcam #ab150115 dilution 1:500). After washing in PBS, the slides were stained with NucBlue Live ReadyProbes Reagent (Invitrogen #R37605). Finally, slides were washed in PBS and mounted with ProLong Diamond Antifade Mountant (Invitrogen, #P36961). Negative controls of primary antibodies omission were carried out in parallel during all processing. NET formation was visualized with the Axio Observer.Z1 inverted microscope equipped with the LSM 880 confocal module (Carl Zeiss, Jena, Germany). All pictures were taken using a 20x/0.5 NA objective. All pictures were prepared using ImageJ (ver. 1.54f, Java 1.8.0_322; 64-bit).

Software and data analysis

Final data visualization and analysis was done in GrapPad Prism 9.1.1 (Dotmatics). For calculating the K_D_ and Ca_0.5_ values a custom modification of specific binding with Hill slope equation was used:

$$Y=\frac{Y_{max}\times X^{h}}{(Z^{h}+X^{h})}+Y_{min}$$

Where Y_max_ is maximal activity from fit, X is a concentration of either GAG or Ca^2+^, h is Hill coefficient, Y_min_ is minimum fitted activity and Z is either K_D_ or Ca_0.5_, depending on the experiment type.

Statistical analysis was done in the same software using ANOVA and Dunnett or Tukey multiple comparison correction as indicated in figure description.
